# Invasive mammals disrupt native dung beetle community coexistence

**DOI:** 10.1101/2024.10.12.617989

**Authors:** Ryo Akashi, Ryo Yamaguchi, Shinji Nakaoka

**Affiliations:** Graduate School of Life Science,Hokkaido University, Sapporo, Hokkaido, Japan; Department of Advanced Transdisciplinary Sciences, Hokkaido University, Sapporo, Hokkaido, Japan

## Abstract

Biological invasions are among the major drivers of biodiversity and are increasing worldwide. Among the invasive species, mammals have a particularly profound impact on native ecosystems. As primary decomposers of mammalian feces, dung beetles are critical in ecosystem functioning, and their community structure is closely linked to their services. However, the introduction of invasive mammals threatens these beetles and potentially disrupts their ecosystem services. Therefore, it is necessary to understand the resulting changes in communities. We developed a novel population dynamics model focusing on the interactions among mammals, feces, and dung beetles. Our results indicate that such invasions increase the risk of extinction of specialist dung beetles that cannot utilize the feces of invasive mammals. The risk of extinction is particularly high when generalist dung beetles show a preference for native feces, leading to intensified interspecific competition for resources. Additionally, the extinction risk of specialist dung beetles increases when invasive mammals display irruptive population dynamics. In conclusion, our findings demonstrated that non-native mammalian invasions disrupt the coexistence of native dung beetle communities, potentially leading to losses in biodiversity and ecosystem function. These risks should be considered in future empirical studies to evaluate the impact of invasive mammals on dung beetle communities.

## 1 Introduction

In the Anthropocene era, the sixth mass extinction represented a profound threat to global biodiversity, resulting in an unprecedented ecological crisis (Pievani 2014; Turvey and Crees 2019). Erosion of biodiversity is directly linked to a decline in ecosystem functions and services, which are essential for environmental stability (Cardinale et al. 2012). Among the primary drivers of biodiversity loss, biological invasions are particularly sequential (Bellard et al. 2016). Invasive species affect native ecosystems through many interspecific interactions, including predation, competition, hybridization, and the introduction of pathogens or parasites (Pyšek and Richardson 2010). These interactions often lead to significant alterations in the community structure, such as reductions in species richness, defaunation, and extinction of native species, as well as extensive ecosystem modifications (Duenas et al. 2021). Therefore, biological invasion is key to maintaining biodiversity and ecosystem conservation. Furthermore, the economic costs associated with biological invasions are escalating, further complicating efforts to achieve ecological and economic sustainability (Diagne et al. 2021).

Conversely, invasive species can sometimes fulfill beneficial ecological roles, effectively substituting native species in certain contexts. For example, invasive species can replace native prey species (Novaro, Funes, and Walker 2000), enhance the redundancy of ecological functions (Onodera, H. S. Enari, and H. Enari 2022), and maintain their diversity as top predators (Letnic et al. 2009). Intentionally introducing non-native species is often undertaken to provide specific ecological services. (e.g., bees as pollinators (Seitz, Vanengelsdorp, and Leonhardt 2020), dung beetles as dung decomposers (Pokhrel, Cairns, Hemmings, et al. 2021), hunting animals for rajor hunting (Carpio et al. 2017), and fish for aquaculture (Gozlan et al. 2010)). From this perspective, non-native species do not always negatively affect native biodiversity and ecological functions. Despite a growing body of case studies on the ecological consequences of communities composed of both invasive and native species, a general understanding of these dynamics needs to be explored.

Non-native mammals have a more significant effect on native ecosystems than other taxa, primarily because of their high trophic levels (Kumschick et al. 2015; Duenas et al. 2021). In particular, they can dramatically alter the amount of mammalian fecal matter in an ecosystem, both through their own defecation and through the reduction of fecal matter from native mammals owing to competitive displacement by non-native species.

Dung beetles are important when considering the effects of non-native mammals on native ecosystems. As primary decomposers of mammalian feces, dung beetles are excellent bioindicators because of their sensitivity to environmental changes and ease of study (A. J. Davis et al. 2001; Bicknell et al. 2014). They are key in terrestrial ecosystems, particularly in forests and pastures, by providing numerous ecological functions beyond the decomposition of feces and carrion. These functions include nutrient cycling, bioturbation, plant growth enhancement, secondary seed dispersal, and fly control (E. Nichols et al. 2008; deCastro-Arrazola et al. 2023). Notably, their community structure, primarily functional diversity, affects ecosystem functions and services (Milotíc et al. 2019; Jorge Ari Noriega et al. 2023).

Community ecologists have conducted both empirical and theoretical studies to understand biodiversity, particularly in the context of community assembly, dynamics, and structure. This contributed significantly to our understanding of ecosystems (May and Arthur 1972; Armstrong and McGehee 1976; Chesson 2000; Tilman 2004; Mougi and Kondoh 2012). Theoretical studies have also been pivotal in exploring the effects of invasive species on native communities (Courchamp, Langlais, and Sugihara 1999; Takimoto and Nishijima 2022). However, research on dung beetle communities often yields conflicting or context-specific results (Errouissi et al. 2004) because different studies have focused on varying functional groups (e.g., body size, food habitat, reproductive behavior, and circadian rhythm) (Slade et al. 2007; Pryke, Roets, and Samways 2022; Jorge Ari Noriega et al. 2023) and each study site has a unique species composition and community structure of mammals and dung beetles (Bogoni et al. 2016). We developed population dynamics models that considered mammals as resource suppliers, feces as resources, and dung beetles as resource consumers to address this issue. These models enable quantitative evaluation of the effects of invasive mammals on native dung beetle communities, as mathematical models can simulate varying strengths of invasion, fecal preferences of dung beetles, and specific species compositions.

In this study, we first focused on simplified scenarios in which a non-native mammal invaded a native community and examined how this affected the population dynamics of native dung beetles. First, we developed a simple model in which we varied the environmental suitability for invasive mammals and the competition coefficients between the dung beetles. Next, we explored the functional response of generalist dung beetles, that they exhibit four different fecal preferences in the presence of invasive mammals, which displayed unique population dynamics, while the strength of interspecific competition for resources varied. Finally, we incorporated a time-delay into the reproduction of invasive mammals to investigate the influence of this additional complexity on the system. These scenarios were analyzed using mammal–feces–dung beetle population dynamics models. Using these models, we assessed the impact of non-native mammal invasions on the population dynamics and extinction risk of native dung beetle communities.

Population size and the presence of mammals greatly influence dung beetle communities by providing feces as a resource and modifying habitats (Bogoni et al. 2016; Raine and Slade 2019). Dung beetles exhibit generalist feeding habitats (Hanski and Cambefort 2014). However feces produced by non-native mammals are sometimes not used by native dung beetles because of their adaptive evolution to native feces (Bornemissza 1979; Amézquita and Favila 2010). Therefore, the degradation rate of livestock is often low because of the low preference for livestock feces (Amézquita and Favila 2010). Non-native dung beetles have often been introduced to solve this problem (Pokhrel, Cairns, and Andrew 2020). Many dung beetles have defaunated in Madagaskar, contrastly dung beetles can use cattle feces to expand their distribution (Hanski, Koivulehto, et al. 2007; Hanski, Wirta, et al. 2008). However, native dung beetles prefer non-native feces (Onodera, H. S. Enari, and H. Enari 2022). Accordingly, native dung beetles exhibit various preferences for non-native feces. Thus, when assessing the effect of invasive mammals on population dynamics and the coexistence of native dung beetle communities, various preferences or availabilities are needed.

Large herbivores have a more significant effect on dung beetle communities than other mammals (Bogoni et al. 2016; Pryke, Roets, and Samways 2022). They are often introduced for livestock or hunting purposes (Carpio et al. 2017). These herbivores frequently exhibit irruptive dynamics, characterized by rapid population increases, subsequent crashes, and eventual stabilization at a carrying capacity lower than the peak (Leopold 1943; Forsyth and Caley 2006; White, Bruggeman, and Garrott 2007). Such irruptive dynamics can profoundly affect the environment (Kaji, Koizumi, and Ohtaishi 1988). However, the impact of these dynamics on dung beetle communities has not been extensively studied. The reproductive characteristics of non-native mammals may significantly affect the community structure and extinction risk of native dung beetle populations.

## 2 Methods

In this study, we envisioned a scenario in which a community of native mammals and dung beetles coexist but faces the invasion of non-native mammals(fig 1). The presence of invasive mammals introduces direct and indirect competition for food and space between native and invasive mammalian species. Additionally, we classified dung beetles into two types in the focal community, i.e., generalists and specialists, which exhibit distinct responses to invasive mammalian feces. While fecal generalists can utilize both native and invasive mammalian feces as food sources, fecal specialists are limited to utilizing only native mammalian feces. Several studies have reported these foraging strategies (ex. Amézquita and Favila 2010; Hanski, Wirta, et al. 2008). Competition for feces can be modeled as a resource competition process in which the resource (i.e., feces) is consumed by both types of beetles, reducing resource availability over time. In the following section, we introduce a simple mathematical model for the aforementioned population dynamics and describe several extensions of this model that reflect realistic scenarios.

**Figure 1:**
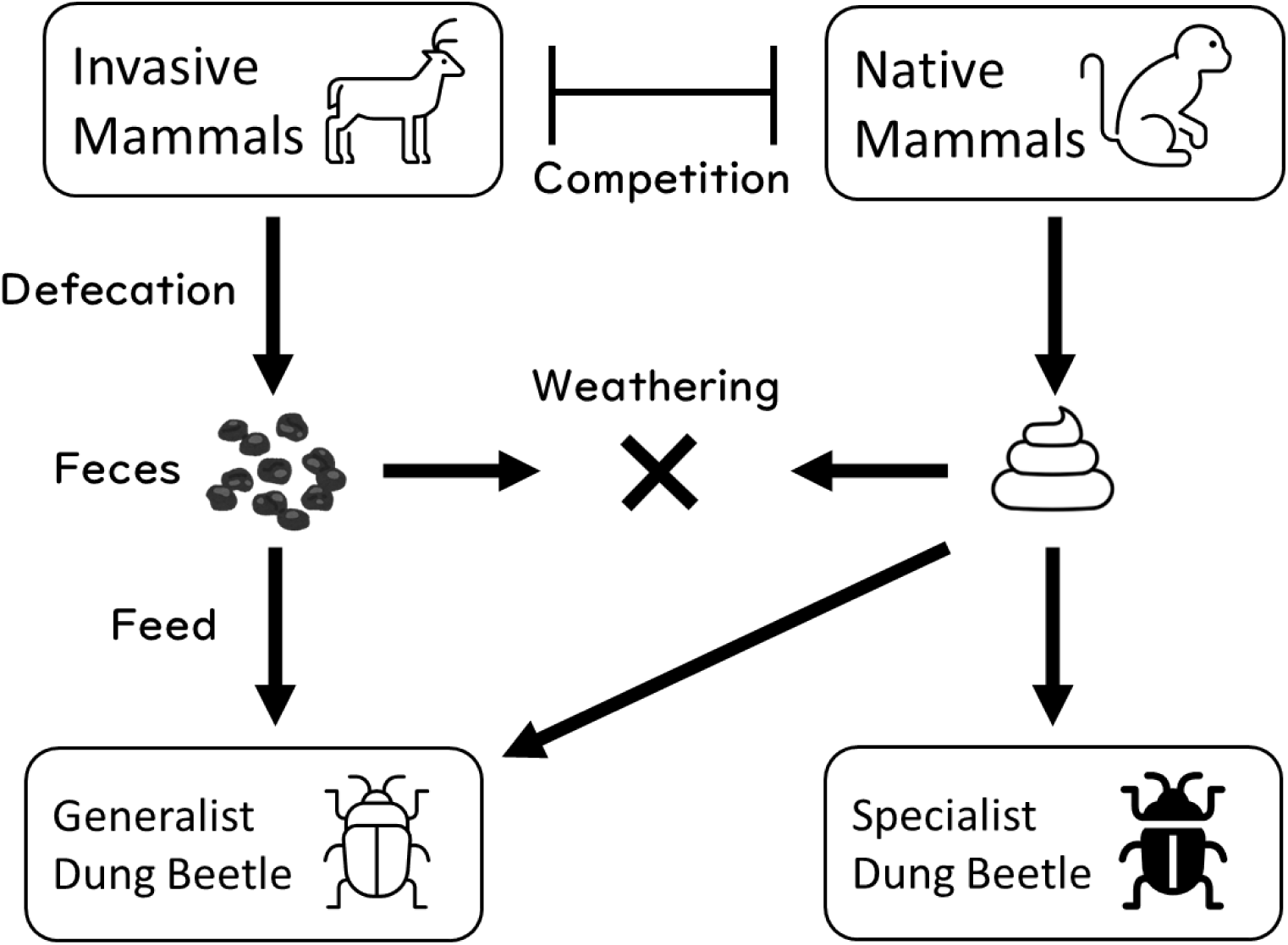
Diagram of the model. Mammals defecate feces. Feces are used by dung beetles. Generalist dung beetles use all feces, but specialists use only native feces.

### 2.1 Model

The population dynamics models used in this study for both mammals and dung beetles were based on the Lotka-Volterra model (Wangersky 1978). *n*2 and *n*1 denote the densities of the native and invasive mammals (individuals per unit area), respectively. The two types of dung beetles are represented by *x* and *y*, where *x* denotes the density of dung generalists and *y* denotes the density of dung specialists. When only a single mammalian species is present, the population dynamics model of mammals takes the form of logistic growth and reaches the carrying capacity (*K*_1_ for *n*_1_ and *K*_2_ for *n*_2_). *r_i_* represents the intrinsic growth rate of each mammal, and *a* is the competition coefficient between the native and invasive mammalian populations. These basic assumptions yield:

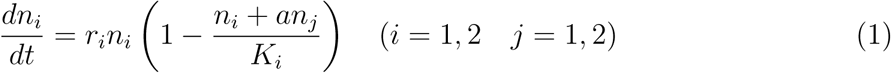

The variable *f_i_*(*i* = 1, 2) is the density of the feces of each mammal, *f*_1_ and *f*_2_ represent invasive and native mammals, respectively. Feces are produced in proportion to mammal abundance and are assumed to weather naturally in proportion to their self-density. *b_i_*(*i* = 1, 2) and *c* are proportionally constant during defecation and weathering, respectively. Feces are consumed by the dung beetles, depending on their density(*x* and *y*) and fecal preference(*F_i_*(*i* = 1, 2), *G*_2_). *F_i_* is the fecal preference of the generalist dung beetle and *G*_2_ is the fecal preference of the specialist. The models of fecal dynamics are written as Eq.2 and 3.

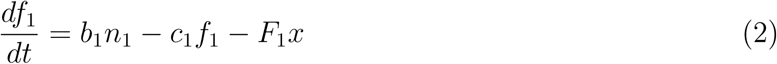

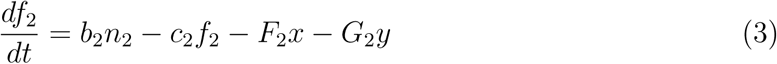

Dung beetle populations exhibit intrinsic growth towards their carrying capacity, which is determined by the availability of mammalian feces. They use feces for breeding (Halffter and Matthews 1966); therefore, their carrying capacity is directly proportional to the amount of available feces. The transformation efficiency from feces to dung beetle reproduction is represented by *e_i_, e_y_*. *h* denotes the competition coefficient for the dung beetles. Generalist dung beetles opportunistically exploit both invasive (*f*_1_) and native (*f*_2_) feces (first and second parts of Eq. 4, respectively). In contrast, specialist dung beetles rely solely on native feces, and their population dynamics follow a classic logistic growth model (Eq. 5).

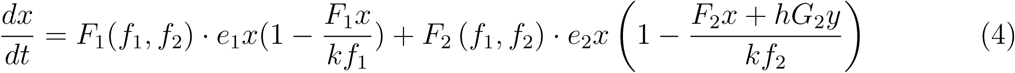

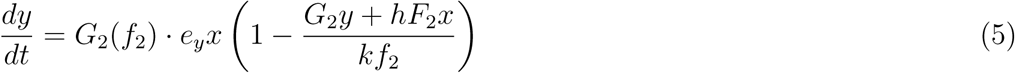

Dung beetle feeding preferences may shift in response to fecal abundance (Tsuji et al. 2021). We adopted the Holling functional response model developed by Leeuwen (Leeuwen, Jansen, and Bright 2007) to represent these dynamics. This model incorporates attack rates (*g_ij_, g_y_*) and handling times (*T_ij_, T_y_*), where *i* denotes the most recently attacked fecal type and *j* represents the previously attacked type. Specialist dung beetles are characterized by attack rate *g_y_* and handling time *T_y_*. Generalists exhibited a Holling Type III functional response, whereas specialists (both pre- and post-invasion) maintained a Holling Type II response following an invasion event.

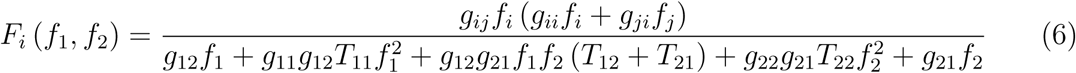

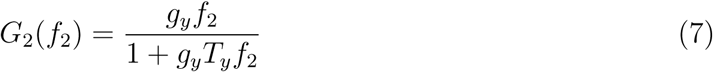

We used these models as scenarios in a numerical analysis to investigate the effects of non-native mammal invasion on native dung beetle communities. Our simulations shed light on the potential consequences of these invasions on the structures and functions of these communities.

**a) Simple model: linear response** When specialists and generalists exhibit identical preferences for native mammal feces(*F*_2_ = *G_y_*) and if we set *g_ii_* = *g_ij_, T_ii_* = *T_ij_* = 1, generalists lose memory of past feces encounters, and handling time is constant. Thus, the fecal preference after invasion *F_i_*(*f*_1_, *f*_2_) is

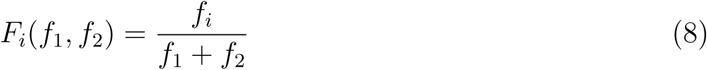

Using eq.1-5,7,8, we simulated the effects of invasive mammals on dung beetle communities. In this study, we investigated the effects of varying the following parameters: *K*_1_, which represents the suitability of the invaded area for invasive mammals, and *h*, which denotes the strength of interspecific competition among dung beetles.

**b) Functional response of generalist’s fecal preference** Previous research has indicated that even generalist feeders may exhibit specific types of functional responses(Abrams 2022). Building on this, we investigated how the different functional responses of generalist dung beetles to mammalian feces impacted specialist populations. We considered four distinct scenarios of generalist fecal preferences (Fig. 2). Preference A (linear response): Generalists utilize both fecal types equally (Eq. 9). Preference B (switching behavior): Generalists strongly favor one fecal type and readily switch when the fecal type becomes scarce. Preference C (high native preference): Generalists strongly prefer native feces but utilize invasive feces. Preference D (high invasive preference): Generalists strongly prefer invasive feces but utilize native feces. The parameter values for each preference scenario are as follows: [*g*_11_, *g*_12_, *g*_21_, *g*_22_] = [[1, 1, 1, 1], [10, 10, 0.1, 0.1], [3, 0.3, 0.9, 0.9], [0.3, 3, 0.9, 0.9]], obtained from (Leeuwen, Jansen, and Bright 2007).

**Figure 2:**
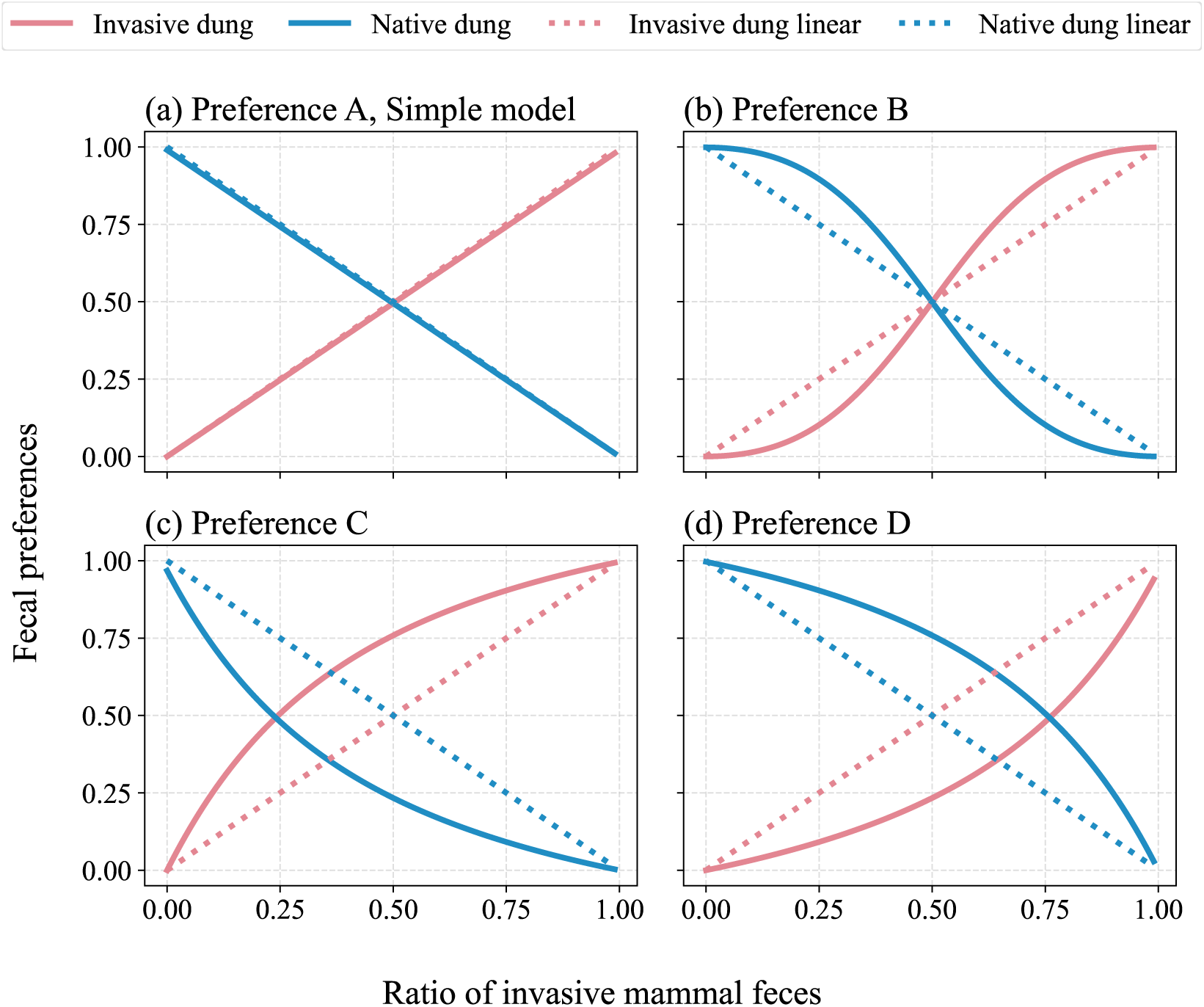
Four scenarios of generalist dung beetle functional response to varying ratios of invasive and native mammal feces. Solid lines depict calculated preferences, while the dotted line represents a linear response (Preference A) for comparison.

**c) Time-delay reproduction of invasive mammals** Large herbivorous mammals can exhibit irruptive population dynamics characterized by rapid increases exceeding their carrying capacity, followed by crashes to lower abundances(Leopold 1943; Scheffer 1951; Forsyth and Caley 2006). Such dynamics are often modeled using time-delay models(Forsyth and Caley 2006). Here, we extend the invasive mammalian population model to a delayed logistic growth model(eq.9) (Hutchinson et al. 1948) to capture these irruptive oscillations. These fluctuations in invasive mammal populations can affect other species through interactions, such as defecation and competition. Using this model, we investigated the effects of the irruptive dynamics of invasive mammals on the dynamics and coexistence of native dung beetle communities.

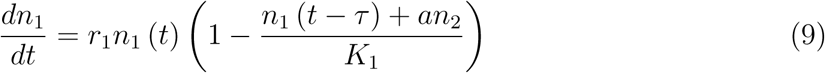

**Table 1:** Notation and parameters used.

| Notation | Description | Value |
| --- | --- | --- |
| $n_1$ | Invasive mammal population density | |
| $n_2$ | Native mammal population density | |
| $x$ | Generalist dung beetle population density | |
| $y$ | Specialist dung beetle population density | |
| $f_1, f_2$ | Density of feces of invasive and native mammals | |
| $r_1, r_2$ | Biotic potential of invasive and native mammals | [0.2, 0.1] |
| $t$ | Time | |
| $K_1$ | Carrying capacity of invasive mammals | [1500, 400-2500] |
| $K_2$ | Carrying capacity of native mammals | [1000] |
| $a$ | Competition coefficient between mammals | [0.4] |
| $b_1, b_2$ | Fecal productivity of invasive and native mammals | [1.0, 1.0] |
| $c_1, c_2$ | Weathering rate of feces | [0.8, 0.8] |
| $F_1, F_2$ | Fecal preference of generalist | |
| $G_2$ | Fecal preference of specialist | |
| $h$ | Competition coefficient between dung beetles | [0-1] |
| $e_1, e_2$ | Invasive and native feces translation efficiency of generalist | [0.1, 0.1] |
| $e_y$ | Native feces translation efficiency of specialist | [0.2] |
| $k$ | Translation efficiency feces to carrying capacity | [1.0] |
| $g_{11}, g_{12}, g_{21}, g_{22}$ | Attack rates of generalist | |
| $g_y$ | Attack rate of specialist | [1] |
| $T_{ij}$ | Handling time of generalist | [1] |
| $T_y$ | Handling time of specialist | [1] |
| $\tau$ | Time-delay of invasive mammals | [0, 6, 8, 10, 12, 14] |

### 2.2 Simulation methods

The numerical analyses were performed using Python (version 3.9.12). Our simulations explored a scenario where non-native mammals invaded a community of coexisting native mammals and two dung beetle species. We constrained the dung beetle competition coefficient 0 ≤ *h* < 1, ensuring that both beetle species were present before the invasion (fig S1). We assumed a higher fecal consumption efficiency for specialists than for generalists based on the greater adaptation of specialist dung beetles to native mammalian feces (*e*_2_ < *e_y_*).

We considered an event in which a small number of non-native mammals(*n*_1_ = 10) invade a community once the population dynamics of the native species reach an equilibrium state. Specifically, we first ran the population dynamics model for 25000 years before the invasion (*t* = 25000), as this duration was sufficient to reach equilibrium (see S1). We then ran the model for an additional 25000 years following the invasion using equations 1-10. If the generalist and specialist population variable amounted to less than the cut-off threshold (1.0 ∗ 10^−6^) during the sumulations, their population dynamics reached an equilibrium state, and the calculation was suspended before 2500 years. The success or failure of mammalian invasion depends on the proportion of invasive to native mammal carrying capacity *K*_1_/*K*_2_, which varies between invasive mammals that failed to invade *K*_1_/*K*_2_ = 0.4 and native mammals that become extinct *K*_1_/*K*_2_ = 2.5. We calculated the ratio of the reduction in specialist dung beetle population density before and after invasion to discuss the extinction risk of the focal community ((*y ∗ −y*)/*y∗*).

When analyzing the time-delay model, we tracked the minimum specialist population density after invasion *y_min_* as density fluctuated. Owing to the oscillations inherent in the model, we set an extinction threshold of 0.1; populations falling below this threshold were considered extinct (assigned a density of 0). The simulations were suspended if *y_min_* remained unchanged for 1000 years. We varied the competition coefficient (*h*) and performed a functional response analysis. This enabled us to investigate how the population dynamics of native and invasive mammals affect the competitive interactions between the two dung beetle populations.

## 3 Result

### 3.1 a) Simple model: linear response

We investigated the effects of invasive mammals on native dung beetle population dynamics using a simplified model. After the invasion, invasive mammal populations stabilized, generalist dung beetles increased, and specialists declined (Fig. S2). We systematically varied the parameters to explore how the carrying capacity of invasive mammals (*K*_1_) and interspecific competition between dung beetles (*h*) influenced the degree of specialist decline. Our model revealed that both the high carrying capacity of invasive mammals (*K*_1_) and the high competition coefficient between dung beetles(*h*) exacerbated the post-invasion decline in specialist dung beetles (Fig. 3a). Conversely, the population of generalist dung beetles and total dung beetle population increased(Fig. S4a, S5a).

**Figure 3:**
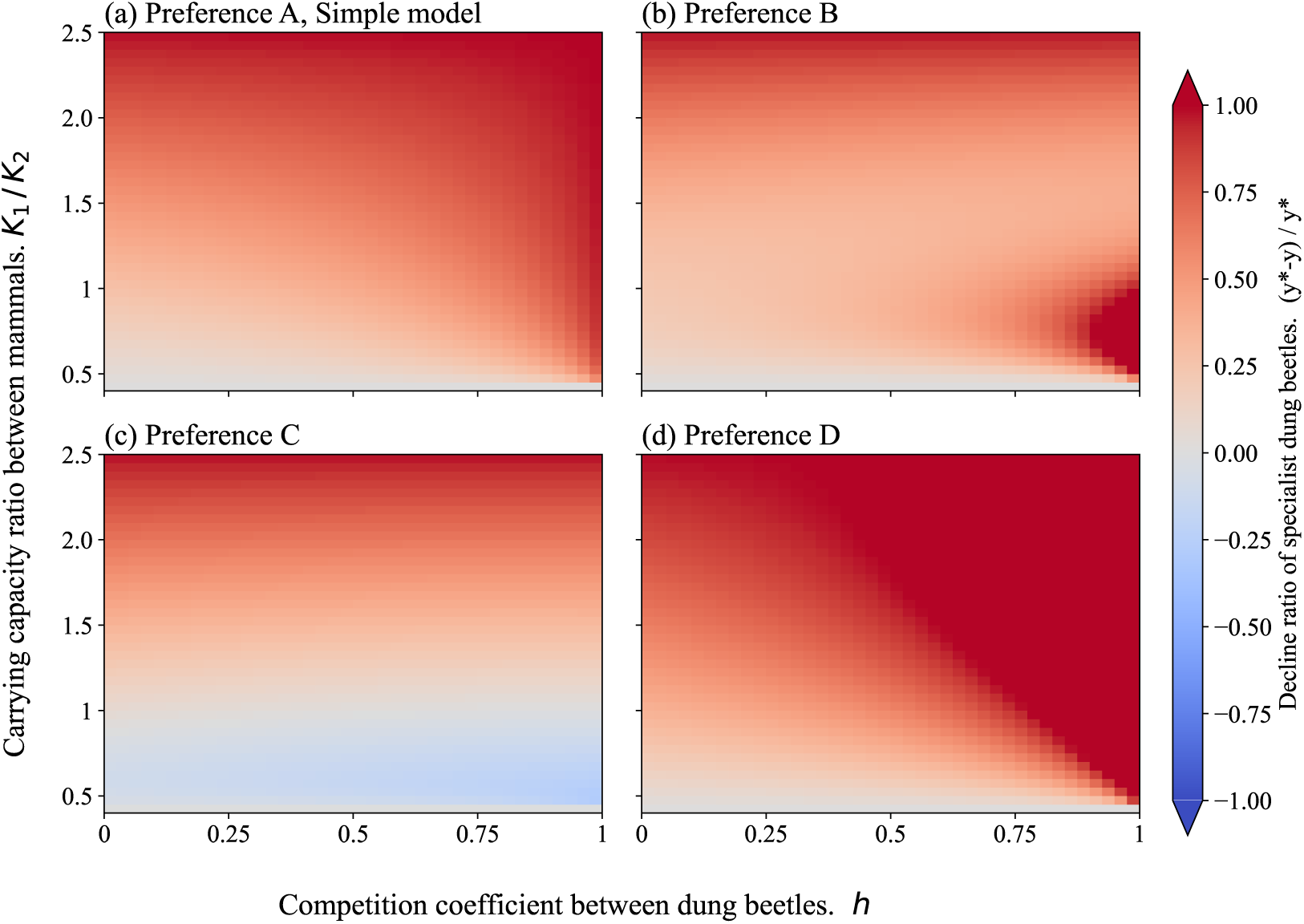
Decline ratio of specialist dung beetles prior to mammal invasion under different fecal preferences. When *K*_1_/*K*_2_ = 0.4, invasive mammals failed to establish; when *K*_1_/*K*_2_ = 2.5, native mammals were competitively excluded. Red plots indicate that the specialist population has decreased prior to the invasion; blue plots indicate that the population has increased. Panels (a), (b), (c), and (d) correspond to Preference A, Preference B, Preference C, and Preference D of generalist dung beetles, respectively.

### 3.2 b) Functional response of generalist’s fecal preference

Different scenarios with generalist fecal preference showed varying declines of specialist dung beetles (Fig. 3). When generalists had no preference between native and invasive feces (Preference A) or preferred native feces (Preference D), an increase in the competition coefficient of dung beetles and the carrying capacity of invasive mammals caused the specialist decline ratio to increase significantly (Fig. 3a, 3d). Preference D caused a stronger decline in specialists than Preference A; high competition coefficients and high carrying capacities of invasive mammals caused the extinction of specialists ((*y^∗^ − y*)/*y^∗^* = 1)(Fig. 3d, S6d).

When generalists preferred invasive feces (Preference C), the specialist decline ratio decreased slightly, and high competition coefficients mitigated the decline in specialists (Fig. 3c). The specialist population increased after the invasion only when the carrying capacity of the invasive mammals was very low 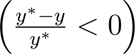 (Fig. S3, S6c). Specialist decline also switched with the carrying capacity of invasive mammals when generalists showed a switching response (Preference B) (Fig. 3b, S6b). Specialists declined strongly with the competition coefficient between dung beetles when the carrying capacity of invasive mammals was low, as under Preference D. However, specialists declined weakly when the carrying capacity was high, as under Preference C.

The trend of specialist decline switching indicated that increasing the carrying capacity of invasive mammals caused a greater decline in specialists when weak competition occurs between dung beetles. However, strong competition caused less decline in specialists (Fig S6b, S7a, S7b, S7c). The increase in generalist and total dung beetle populations was not significantly different from that under the general fecal preference(Fig. S4, S5).

The mechanisms of specialist decline include resource reduction owing to native mammal decline and resource competition between generalist dung beetles. The effect of resource reduction was represented by the declining ratio of specialists when the competition coefficient between dung beetles was zero (*h* = 0) (Fig. S6). We examined intraspecific and interspecific competition strength changes before and after the invasion to reveal the effect of competition on the specialist population. The competition strength changes that species *i* received from species *j* are denoted *C_ij_* (*i, j*) = (*x, x*), (*x, y*), (*y, x*), (*y, y*). From Eq. 4, generalists intraspecific competition is 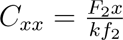, specialists intraspecific competition is 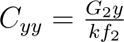, interspecific competition that generalists received from specialists is 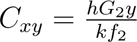, and interspecific competition that specialists received from generalists is 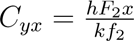. Interspecific and intraspecific competition strengths changed with the carrying capacity of the invasive mammals (Fig. S8). The interspecific competition strength received by specialists increased with the carrying capacity of invasive mammals under Preferences A and D, decreased under Preference C, and showed a switching pattern of increasing and declining with carrying capacity under Preference B.

Interspecific and intraspecific competition changed with the carrying capacity of the invasive mammals and the competition coefficient between dung beetles (Fig. 4, S9, S10, S11). Patterns that decreased the ratio of specialists to interspecific competition specialists were strongly related. Strong competition caused a greater decline, whereas weak competition mitigated the decline of specialists. Under Preference B, the shape of the change in the interspecific competition strength experienced by specialists with increasing competition coefficients was almost the same as that of the declining ratio in the specialist population size (Fig. S6b, S7d).

**Figure 4:**
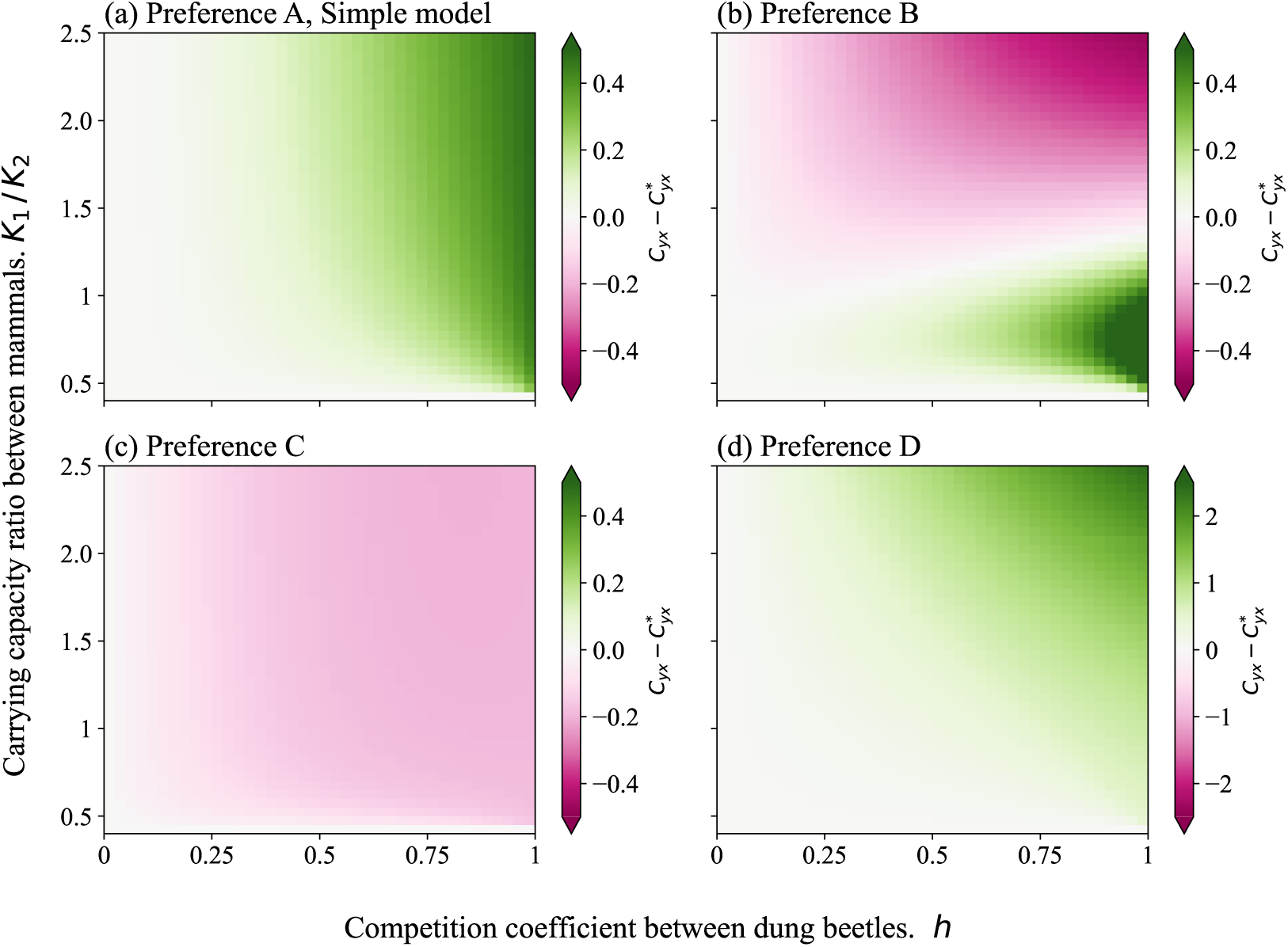
Interspecific competition strength received by specialists 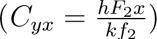 change prior to the invasion. Green plots indicate that competition has increased since before the invasion; purple plots indicate that competition has decreased. Panels (a), (b), (c), and (d) correspond to Preferences A, B, C, and D, of generalist dung beetles, respectively.

### 3.3 c) Time-delay reproduction of invasive mammals

We investigated the effect of the irruptive dynamics of invasive mammals on specialist dung beetles. A large reproductive time-delay caused a greater decline in specialists, increasing their extinction risk (Fig. 5). Excessive competition induced a greater decline of specialists in time-delayed situations, similar to situations without a time-delay. We analyzed the cut-off threshold and found that changing it (0.1 to 1 or 0.01) yielded almost the same results. If generalists have different preferences, the irruptive dynamics of invasive mammals also cause the number of specialists to decrease (Fig. S12).

**Figure 5:**
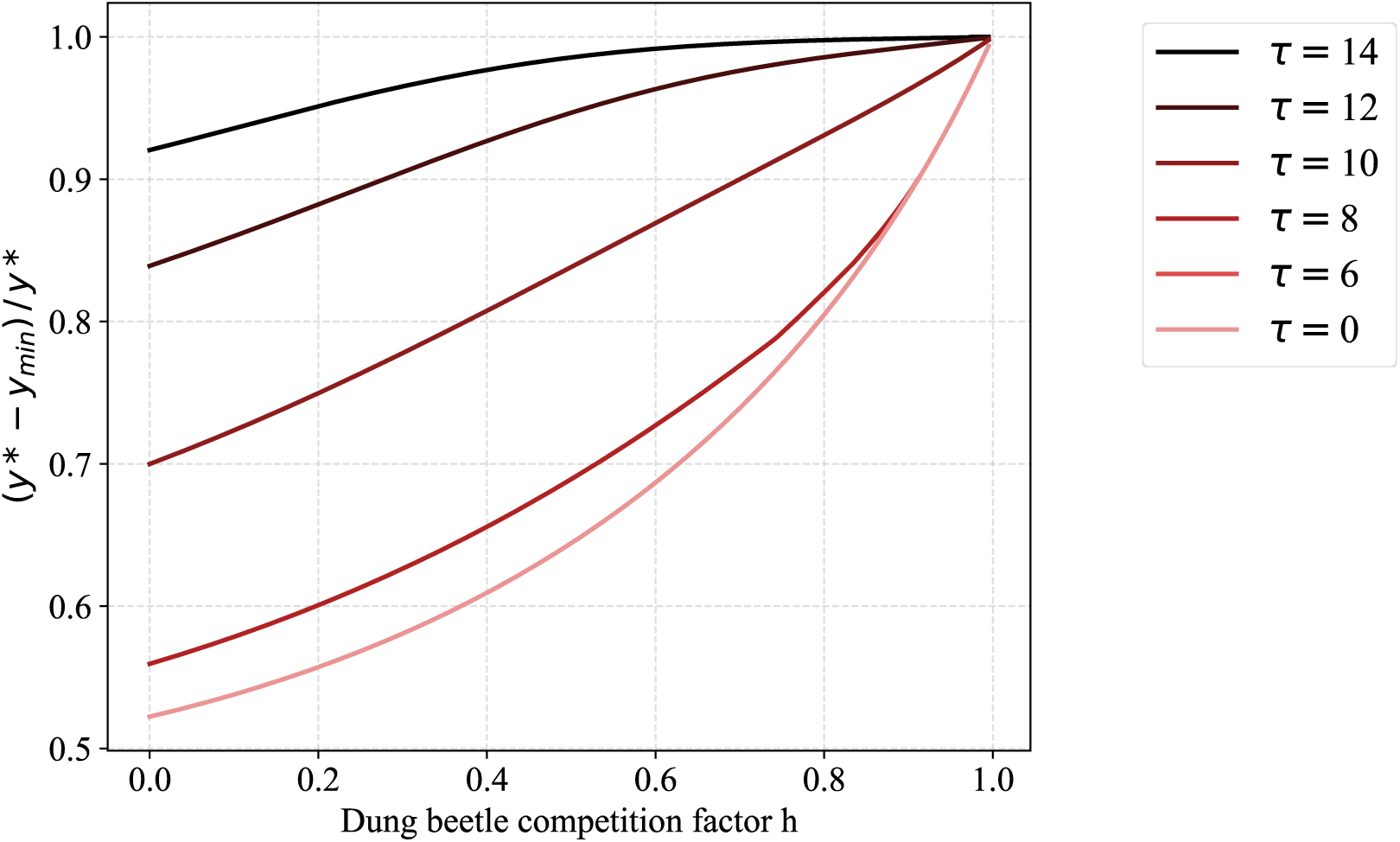
Decline ratio of specialist dung beetles prior to mammal invasion when invasive mammals exhibit irruptive dynamics. The x-axis represents the competition coefficient of dung beetles (*h*), and the y-axis represents the decline ratio of specialists 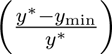. Each line represents a different time-delay (*τ*). The fecal preference of generalists is Preference A, and the carrying capacity of invasive mammals is fixed at *K*_1_ = 1500.

## 4 Discussion

### 4.1 Increase specialist dung beetle extinction risk

Our study demonstrated that the invasion of non-native mammals significantly increases the extinction risk of native specialist dung beetles and decreases the overall coexistence potential of native dung beetle communities. This phenomenon can be explained by two primary mechanisms: first, the competitive displacement of native mammals by non-native mammals reduces the availability of native feces; second, the population increase of generalist dung beetles intensifies the competition for diminished fecal resources.

An increase in the carrying capacity of non-native mammals leads to a decline in native mammalian populations owing to competition for resources, thereby reducing the amount of feces available to specialist dung beetles. The effect of this reduction in fecal resources was quantified using the decline ratio of specialists when the competition coefficient between the dung beetles was zero (h=0) (Fig. 3). The competition intensity experienced by the specialists under a linear Preference A response increased as the interspecific competition coefficient increased (Fig. 4a). Following this invasion, generalist dung beetles could exploit more fecal resources, increasing in the population. This, in turn, results in a decline in specialist dung beetle populations owing to intensified interspecific competition (Fig. 3 and 4).

This study showed that generalist dung beetles have a competitive advantage over specialists. An example of this relationship between generalists and specialists was reported in Madagascar. Hanski, Wirta, et al. 2008 documented that three species of *Helictopleurini* dung beetles, which can utilize non-native cow feces, have expanded their habitat range more widely than other species within the same subgenus. Furthermore, Hanski, Koivulehto, et al. 2007 found that 22 of 51 *Helictopleurini* dung beetle species have become extinct or have experienced population decline. They attributed the defaunation of many dung beetle species to factors such as deforestation, fragmentation of natural habitats, and the historical extinction of large mammals. According to the present study, interspecific competition between generalist and specialist dung beetles may also play a significant role in these declines.

The effects of invasive species that disrupt native food webs are well-documented. For example, competition between non-native and native plants can alter pollinator resource selection (Brown, Mitchell, and Graham 2002; Goodell and Parker 2017), and invasive mammals can replace native prey (Novaro, Funes, and Walker 2000). The concept of this study, which focused on the competition between non-native and native species and the resulting resource selection by native species, is valuable for studying non-native species in other taxa. Terui et al. 2023 found that the intentional release of native species can intensify intraspecific and interspecific competition, thereby undermining the ecological stability of native communities. Overall, this and previous studies showed that the invasion of non-native and native species intensifies competition within native ecosystems, leading to a decline in the diversity and population of native communities. This model, developed for the invasion of non-native mammals, could be applied to range expansion and colonization if both specialist and generalist dung beetles exist. Climate change causes range shifts in native and non-native species (Parmesan 2006), especially Cervidae, which is a range-changing mammal due to climate change (Dawe and Boutin 2016; Weiskopf, Ledee, and Thompson 2019). In the Tohoku region of Japan, the range expansion of native sika deer driven by factors such as global warming and human depopulation poses a growing problem (Ohashi et al. 2016). Additionally, some dung beetles are not attracted to the feces of these deer (Tsuji et al. 2021), raising concerns about the potential decline in dung beetle populations. This decline could be particularly critical, as sika deer populations are known to exhibit irruptive dynamics (Kaji, Koizumi, and Ohtaishi 1988) (further discussed in Section 4.3 Time-delay reproduction of invasive mammals).

### 4.2 Functional response of generalist’s fecal preference

The fecal preference of generalist dung beetles significantly affects the extinction risk of specialist dung beetles (Fig. 3). Interspecific competition intensifies when generalists prefer native feces (Preference D), increasing the extinction risks for specialists (Fig. 3d, 4d). For example, native roller dung beetles that do not utilize invasive cow feces are in the minority in Mexico, whereas native generalists who prefer native monkey feces are in the majority (Amézquita and Favila 2010). This scenario reflects Preference D, raising concerns about further population decline and the potential extinction of roller dung beetles. Roller dung beetles contribute significantly to dung removal and dispersal (Huerta, Arellano, and Cruz 2018; Manns et al. 2020); therefore, their decline would result in the loss of ecosystem function.

Niche partitioning occured when generalists prefer invasive feces (Preference C), which can improve or mitigate the possibility of coexistence (Fig. 3c, 4c). Many native dung beetles prefer invasive civet feces to native raccoon dog feces in Japan (Onodera, H. S. Enari, and H. Enari 2022). In this situation, the impact of invasive mammals on the native dung beetle community would be limited if specialists are present because of reduced resource competition.

Dung beetles are mainly generalist consumers, and some studies have shown they shift their resources (Hanski, Wirta, et al. 2008; Stavert et al. 2014). When generalists exhibit a switching preference (Preference B), the bifurcation of the decline ratio of specialists depended on the carrying capacity of invasive mammals (Fig. 3b, S6b). Consequently, with weak competition among dung beetles, an increase in the carrying capacity of invasive mammals leads to a greater decline in the number of specialists. Conversely, the decline in the number of specialists was less pronounced with strong competition among dung beetles (Fig. S6b, S7a, b). In these models, the effect of switching predation on decreasing extinction risk was limited. The coexistence of dung beetles was only promoted when the population of invasive mammals was large. Previous theoretical and empirical studies have shown that predation switching promotes coexistence (Teramoto, Kawasaki, and Shigesada 1979; Sundell et al. 2003). Mammal feces as a resource are produced in proportion to mammalian populations and are not based on the density of feces. Therefore, the consumption of dung beetles did not affect feces production. This relationship differs from that of ordinarily living prey; as a result, switching predation does not always contribute to the coexistence of dung beetles.

The functional response of various predators has been studied by behavioral experiments with density manipulation (Sundell et al. 2003; Timms et al. 2008). Therefore, the fecal preference of dung beetles has also been studied using display experiments (Mansourian et al. 2016; A. G. Jones et al. 2012) and dung-baited trap sampling (Whipple and Hoback 2012; Tsuji et al. 2021). Previous studies have shown that the fecal detection and selection of dung beetles are caused by receiving volatile organic compounds from feces via olfactory receptors (Inouchi et al. 1987; INOUCHI, SHIBUYA, and HATANAKA 1988; Frank, Brückner, et al. 2018). Except for behavioral experiments, olfactory receptor genomic analysis and electrophysiological studies of olfactory cells (ex. Chen, Li, and Shao 2019) may reveal fecal detection and preference because larval feeding experience did not affect adult olfactory response of dung beetles (Dormont et al. 2010).

### 4.3 Time-delay reproduction of invasive mammals

This study showed that time-delayed reproduction causes a higher extinction risk in native specialist dung beetles (Fig. 5). Irruption or time-delayed reproduction often results in large herbivores (Scheffer 1951). The reasons why irruption and a larger time delay occur are explained by environmental suitability, invasion of new habitats, release from predators and hunting, and delay of mating behavior caused by sociality and age at first reproduction due to feeding conditions associated with increased population density (DR 1997). In conclusion, we need to focus on what species invade and what parameters of population dynamics depend on the environment to assess the quantitative impact on native communities.

Irruption significantly affects the vegetation and soil (Kaji, Koizumi, and Ohtaishi 1988; Starns et al. 2015). Large herbivores affect the dung beetle community through habitat modification (e.g., vegetation cover and soil hardness) (Iida, Soga, and Koike 2018).

Therefore, when considering the impact of mammals on dung beetles, direct interactions, such as supplying feces, and indirect interactions, such as habitat modification without resource interactions, are important. This result may be complicated considering habitat modification by mammals; thus, upgrading the theoretical model to include habitat factors is necessary.

### 4.4 Limitation and Future research question

We did not consider the evolutionary dynamics of dung beetles or mammals in this study. However, evolution is crucial, as both non-native and native species are exposed to novel selective pressures during the process of adaptation by non-native species to the new environment, as well as their interactions with native prey (Lee 2002; Yoshida, Goka, et al. 2007). Evolution, especially rapid evolution, can influence population dynamics and, in some cases, reduce extinction risk through mechanisms such as evolutionary rescue (Yoshida, L. E. Jones, et al. 2003; Bell 2017). If evolutionary dynamics were considered in this study, specialist and generalist dung beetles might have evolved fecal preferences (niche shifts), and mammals could have evolved population dynamic parameters (e.g., growth rate), potentially facilitating coexistence. Unfortunately, studies focusing on the genetic basis of fecal preference, particularly for olfactory receptors, have not been conducted. Therefore, investigating the evolution of fecal preferences using theoretical and empirical approaches remains challenging. Nonetheless, incorporating evolutionary dynamics and expanding the model would be valuable to better understand the interactions between mammals and dung beetles.

Additionally, we did not consider the feedback effects from dung beetles to mammals in our models because of the need for simplification and to focus on separating the mechanisms of decline. However, dung beetles serve as food resources for some carnivorous and omnivorous mammals (RANSOME and HUTSON n.d.), enhance plant growth by relying on herbivores (deCastro-Arrazola et al. 2023), and regulate mammalian parasites (E. Nichols et al. 2008). These interactions suggest that dung beetles provide positive feedback for mammalian populations. Including these feedback effects complicates the dynamics of dung beetles and potentially influences their likelihood of coexistence.

We focused solely on fecal availability and dung beetle selection as mechanisms for niche partitioning and functional grouping but did not consider other factors. Our findings showed that differences in fecal preferences among dung beetles are crucial for their coexistence. However, (Frank, Krell, et al. 2018) pointed out that fecal specialization does not fully explain latitudinal diversity in dung beetles. Other factors such as differentiation in seasonal and day/night activities (Hanski and Cambefort 2014), dung discovery speed (Jacobs et al. 2008), and spatial partitioning into microhabitats (Mehrabi et al. 2014) may also promote coexistence. The relationships between the functional groups of dung beetles and their ecological roles, particularly in fecal removal and decomposition, are well documented. For example, larger dung beetles remove more feces (Nervo et al. 2014; Gebert et al. 2022), and different feeding and reproductive strategies (dwellers, tunnellers, and rollers) use varying amounts of feces (Chen, Li, and Shao 2019). Thus, the diversity and structure of functional groups, including activity patterns, body sizes, and reproductive categories, are important for fecal removal (Slade et al. 2007). Additionally, differences in body size and feeding habitats contribute to the dynamics of dung beetle communities (Whipple and Hoback 2012; Bogoni et al. 2016). In conclusion, both empirical and theoretical studies that consider the functional groups of dung beetles and mammals, along with niche partitioning beyond fecal preference, are necessary to understand the diversity of dung beetle species and the impact of mammals on ecological functions within dung beetle communities better.

## 5 Conclusion

We developed a new population dynamics model that explains the various scenarios in which the dynamics of dung beetle communities are affected by the invasion of non-native mammals. We found that non-native mammal invasions disrupted the coexistence of native dung beetle communities, particularly when different beetle species relied on varying fecal resources. The population dynamics of dung beetles have become more complex, depending on the fecal preferences of other dung beetle species that utilize non-native feces. A higher preference of generalist dung beetles for non-native feces facilitates coexistence, whereas a preference for native feces increases the decline of dung beetle species that cannot utilize non-native feces, making coexistence more difficult. These declines are explained by a reduction in native fecal resources due to competition, as well as interspecific resource competition among dung beetles. When invasive mammals exhibit irruptive population dynamics, they destabilize native dung beetle communities. This study demonstrated that it is essential to consider dung beetle fecal availability and preferences, the carrying capacity of invasive mammals, and the characteristics of invasive mammal population dynamics to assess the impact of invasive mammals on native dung beetles. In conclusion, an increase in invasive mammals may lead to a greater decline in dung beetles with specialized feeding habits.

## Supporting information

Spplimental file

## 6 Acknowledgment

We thank the members of laboratory of mathematical biology and Hayakawa Lab for support, and Dr.Takashi Hayakawa, Dr.Shirow Tatsuzawa, Dr.Hiroaki Sato, Dr.Akari Matsuki, Dr.Akane Hara, Dr.Takuma Niida, Dr.Yoichi Tsuzuki,Dr.Nozomi Akashi, and Mr.Junya Sunagawa for their helpful comments. This work was supported by JST SPRING, Grant Number JPMJSP2119 (to R.A.), JST Grant Numbers JPMJPF2108, JPMJCR23J4, Japan Agency for Medical Research and Development grant number 22gm1710004h0001 (to S.N.).

## 7 Author contributions

R.A.:conceptualization, formal analysis, funding acquisition, investigation, methodology, validation, visualization, writing―original draft, and writing―review and editing; R.Y.:conceptualization, methodology, supervision, writing―review and editing; N.S.:funding acquisition,supervision, writing―review and editing. All authors provided their final approval for publication and agreed to be held accountable for the work performed.

