## Supplementary material for "Invasive mammals disrupt native dung beetle community coexistence": Spplimental file

October 12, 2024

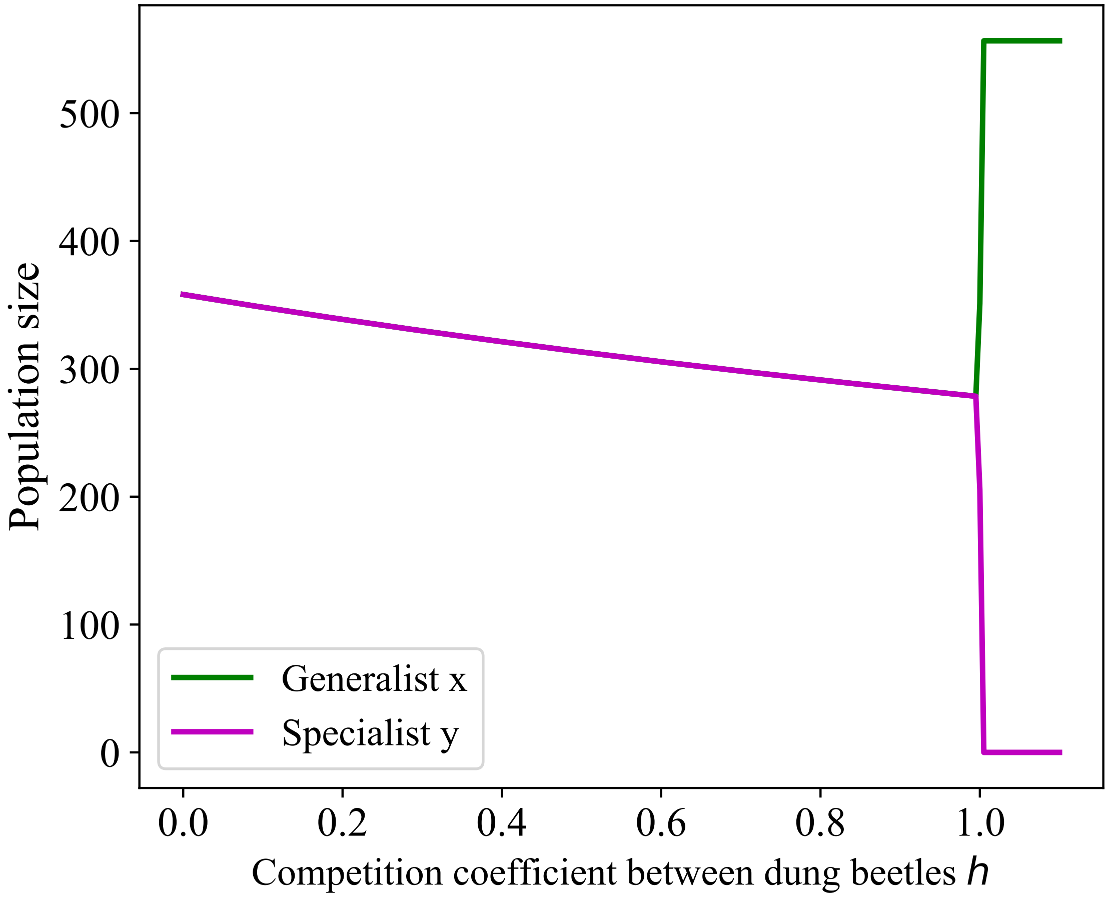


Fig. S1: Population size of dung beetles before mammal invasion with different competition coefficient. Competition coefficients between dung beetles varied 0 to 1.1 in 0.05 increments. Green line represents generalist dung beetles and purple line represents specialist dung beetles.

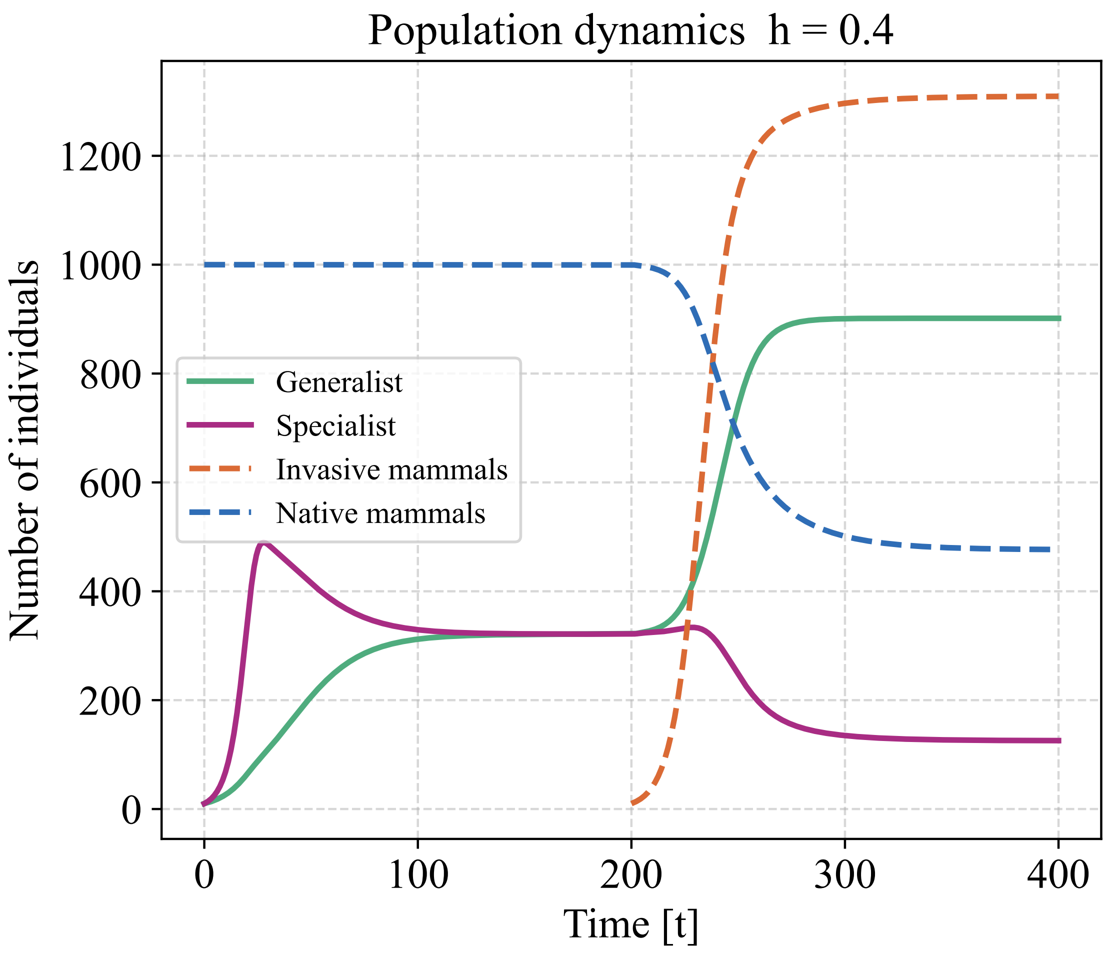


Fig. S2: Population dynamics of mammals and dung beetles. Carrying capacity of invasive mammals($K_1 = 1500$), competiton coefficients of dung beetles ($h = 0.4$), initial population size of dung beetles is $x_0 = y_0 = 10$ and fecal preference of generalist dung beetle is Preference A; no preference shown for feces. Invasive mammals invade native community when t = 200. Blue dot line is native mammal, orange dot line is invasive mammal, green line is generalist dung beetle, purple line is specialist dung beetle.

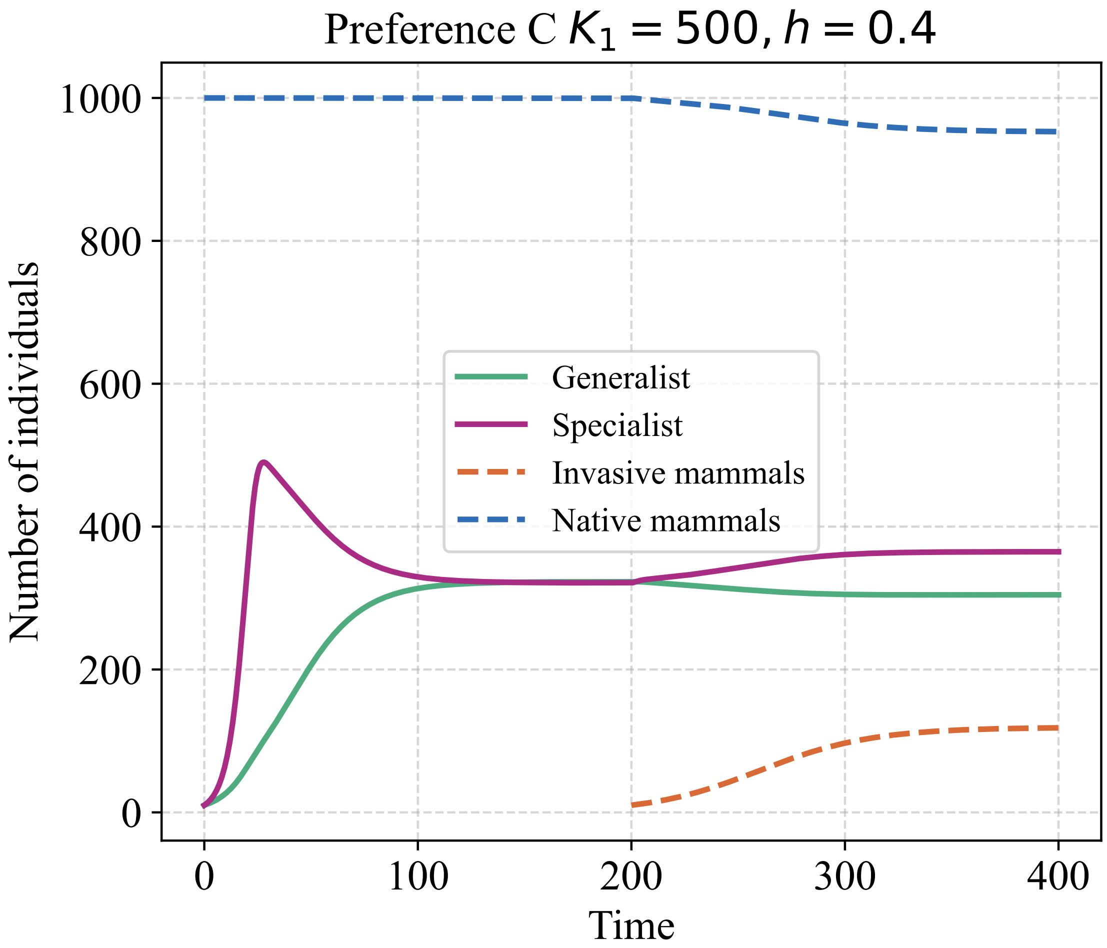


Fig. S3: Population dynamics of mammals and dung beetles when generalist prefer invasive feces and low carrying capacity of invasive mammals. Carrying capacity of invasive mammals($K_1 = 500$), competiton coefficients of dung beetles ($h = 0.4$), initial population size of dung beetles is $x_0 = y_0 = 10$ and fecal preference of generalist dung beetle is Preference C. Invasive mammals invade native community When t = 200. Blue dot line is native mammal, orange dot line is invasive mammal, green line is generalist dung beetle, purple line is specialist dung beetle.

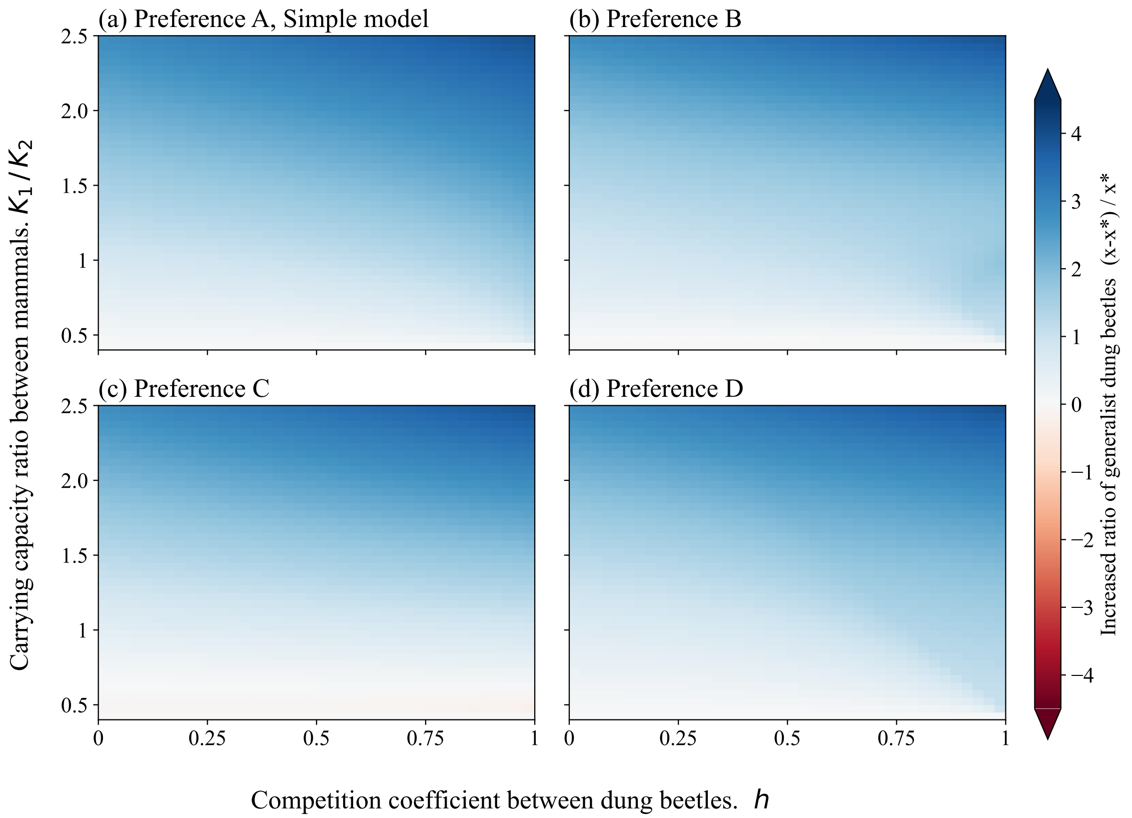


Fig. S4: Increase ratio of generalist dung beetles from before mammal invasion under different fecal preferences. When the plot color is blue, generalist population has increased since before the invasion; when the plot color is red, population has decreased. Panels (a), (b), (c), and (d) correspond to different fecal preferences of generalist dung beetles: Preference A, Preference B, Preference C, and Preference D, respectively.

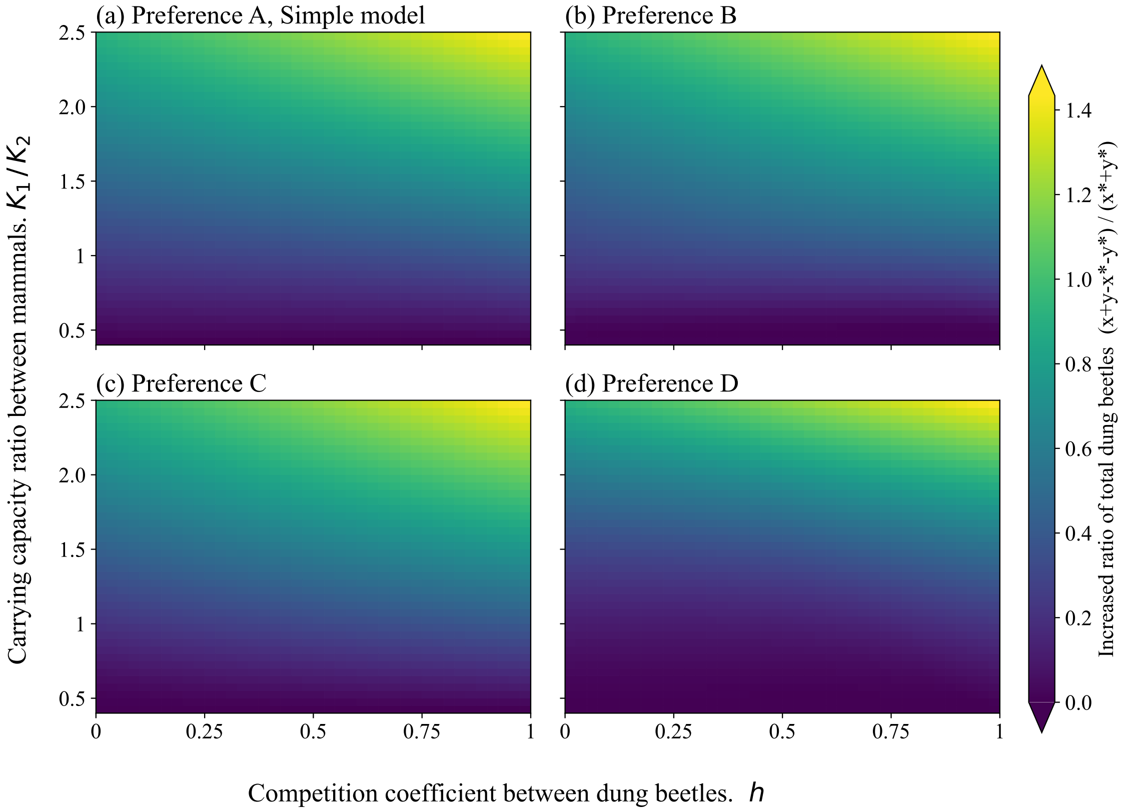


Fig. S5: Increase ratio of total dung beetles from before mammal invasion under different fecal preferences. When the plot color is yellow, generalist population has increased since before the invasion; when the plot color is blue, population has decreased. Panels (a), (b), (c), and (d) correspond to different fecal preferences of generalist dung beetles: Preference A, Preference B, Preference C, and Preference D, respectively.

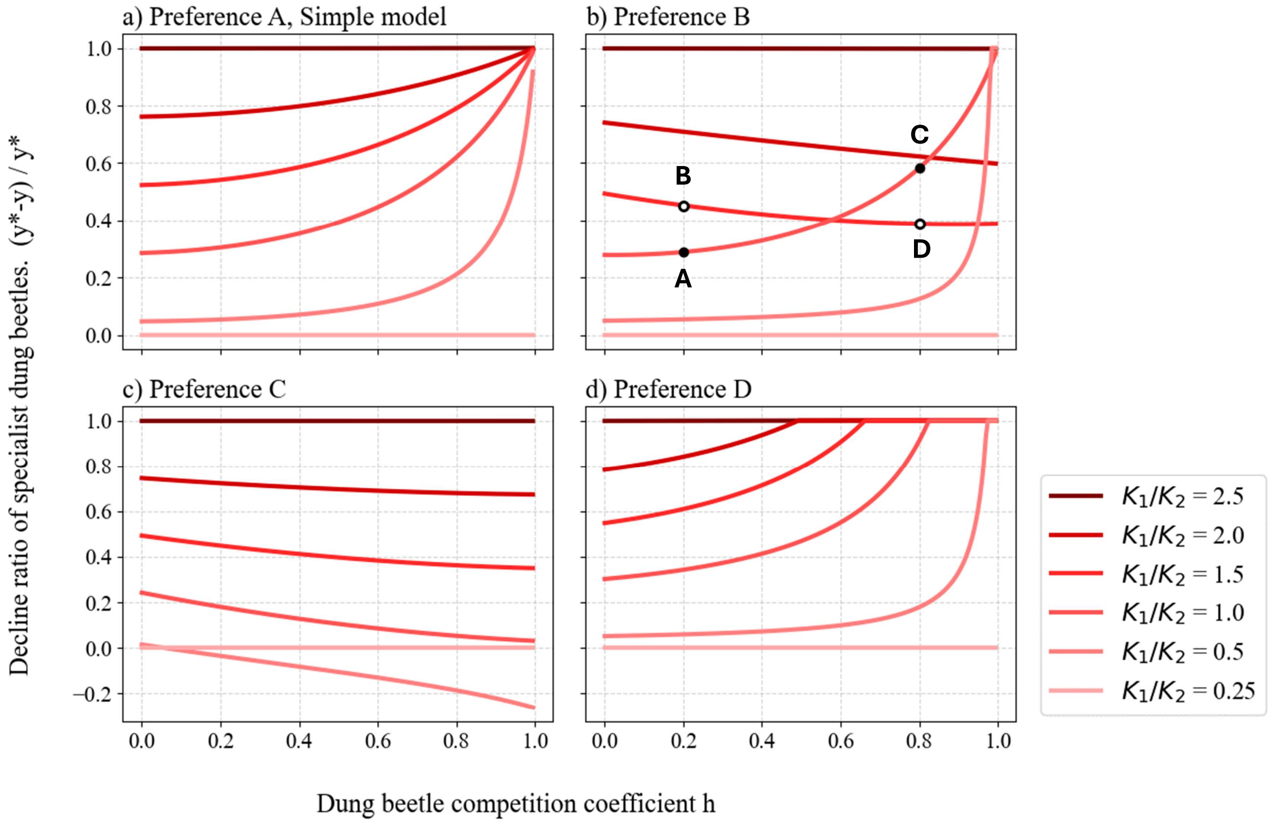


Fig. S6: Decline ratio of specialist dung beetles from before mammal invasion under different dung preferences. X-axis is competition coefficient between dung beetles (h), y-axis is decline ratio of specialist dung beetle from before mammal invasion, and each line in the panels represents a different carrying capacity of invasive mammals. When $K_1 = 250$, invasive mammals failed to establish; when $K_1 = 2500$, native mammals were competitively excluded. Panels (a), (b), (c), and (d) correspond to different fecal preferences of generalist dung beetles: Preference A, Preference B, Preference C, and Preference D, respectively.

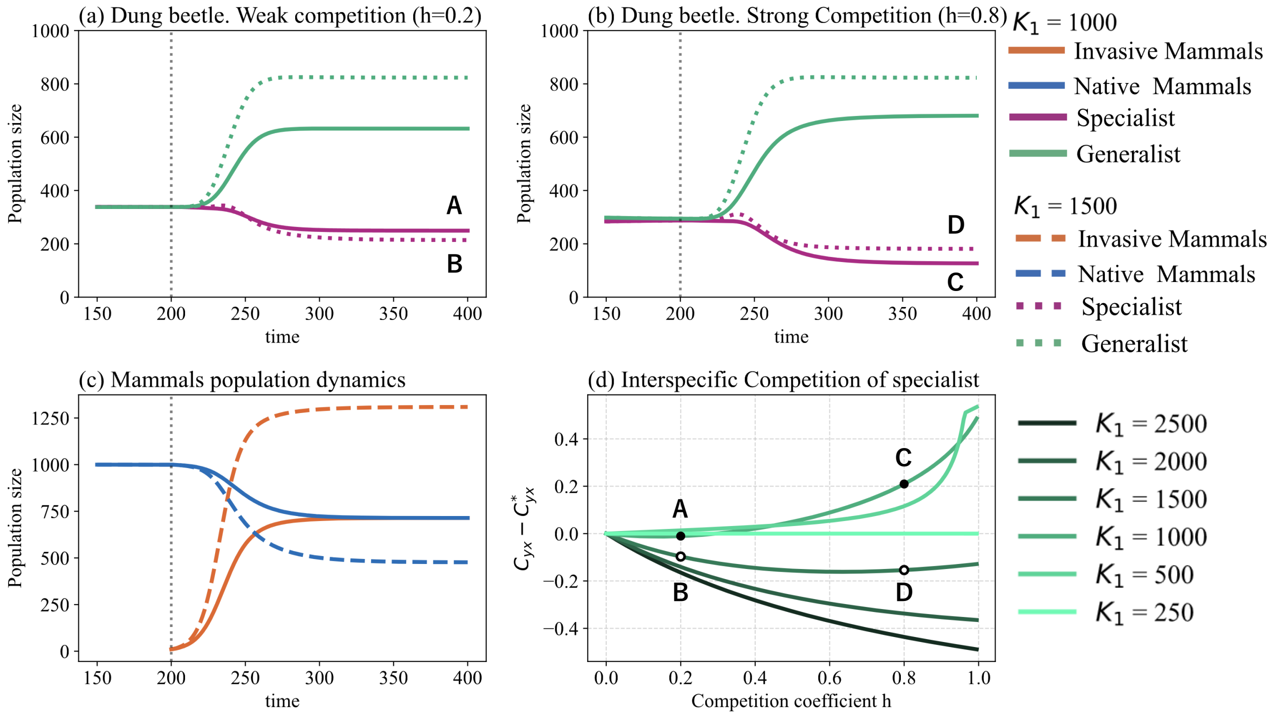


Fig. S7: Population dynamics and interspecific competition strength difference when fecal preference of generalist is Preference B. (a) Population dynamics of dung beetles with weak competition coefficient(h=0.2) between dung beetles. (b) Population dynamics of dung beetles with strong competition coefficient(h=0.8) between dung beetles. Green lines represent generalist dung beetles and purple line represent specialist. (c) Population dynamics of mammals. Blue line represent native mammals and orange line represent invasive mammals. Panels (a), (b) and (c), invasive mammals invade native community when t = 200 and solid lines represent the situation when low carrying capacity of invasive mammals ($K_1 = 1000$) and dot lines represent high caryying capacity($K_1 = 1500$). (d) Interspecific competition strength received by specialist($C_{yx} = \frac{hF_2x}{kf_2}$) difference from before the invasion. Y-axis represents difference of interspecific competition of specialist, if value take positive interspecific competition get intensified since before invasion. Character A, B, C and D mean the same set of parameters.

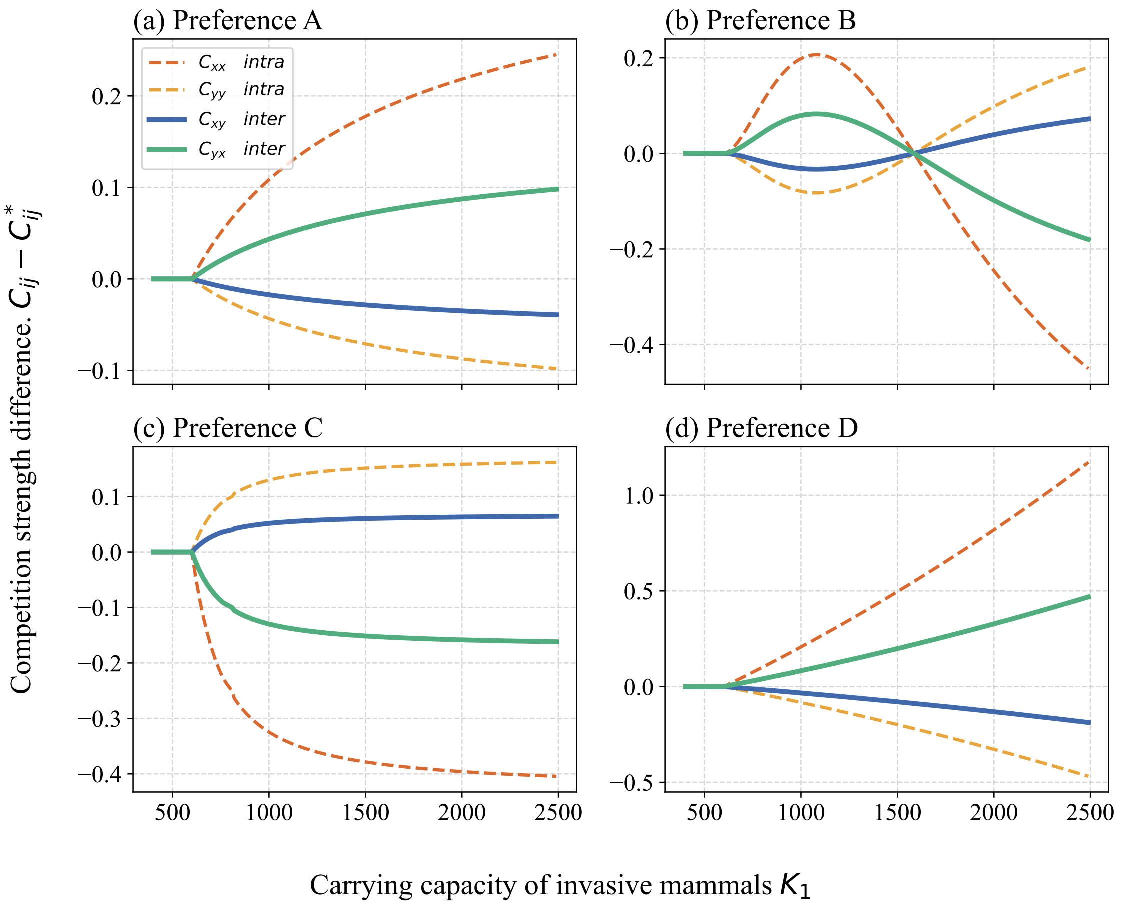


Fig. S8: Difference of competition strength with varying carrying capacity of invasive mammals. Solid lines represent interspecific competition and dot lines represent intraspecific competition. $C_{xx}$, $C_{yy}$, $C_{xy}$, $C_{yx}$ means competition strength: intraspecific competition of generalist, intraspecific competition of specialist, interspecific competition received by generalist and interspecific competition received by specialist respectively. Panels (a), (b), (c), and (d) correspond to different fecal preferences of generalist dung beetles: Preference A, Preference B, Preference C, and Preference D, respectively.

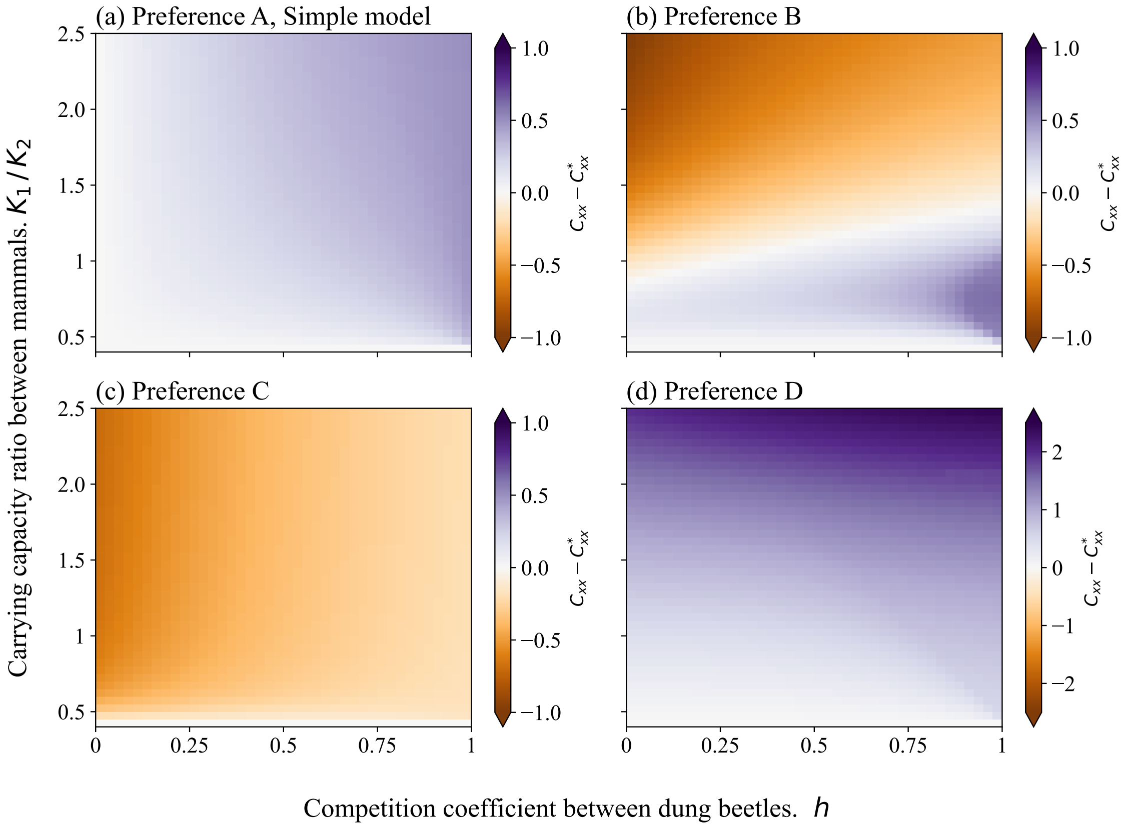


Fig. S9: Intraspecific competition strength of specialists ($C_{xx} = \frac{F_2 y}{k f_2}$) difference from before the invasion. X-axis represents the competition coefficient between dung beetles (h), y-axis represents competition coeffcient of dung beetles. When the plot color is green, competition has increased since before the invasion; when the plot color is brown, competition has decreased. Panels (a), (b), (c), and (d) correspond to different fecal preferences of generalist dung beetles: Preference A, Preference B, Preference C, and Preference D, respectively.

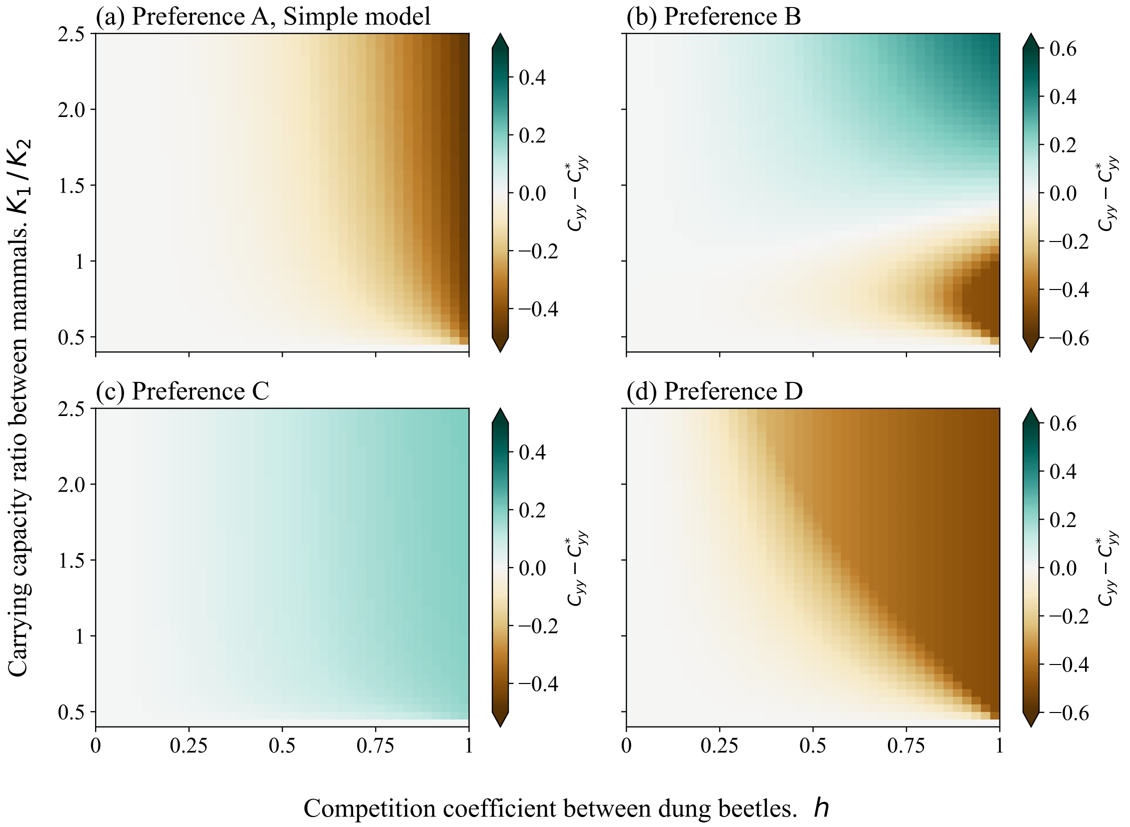


Fig. S10: Intraspecific competition strength of generalsits ($C_{yy} = \frac{G_2 y}{k f_2}$) difference from before the invasion. X-axis is the competition coefficient between dung beetles (h), y-axis represents competition coeffcient of dung beetles. When the plot color is green, competition has increased since before the invasion; when the plot color is brown, competition has decreased. Panels (a), (b), (c), and (d) correspond to different fecal preferences of generalist dung beetles: Preference A, Preference B, Preference C, and Preference D, respectively.

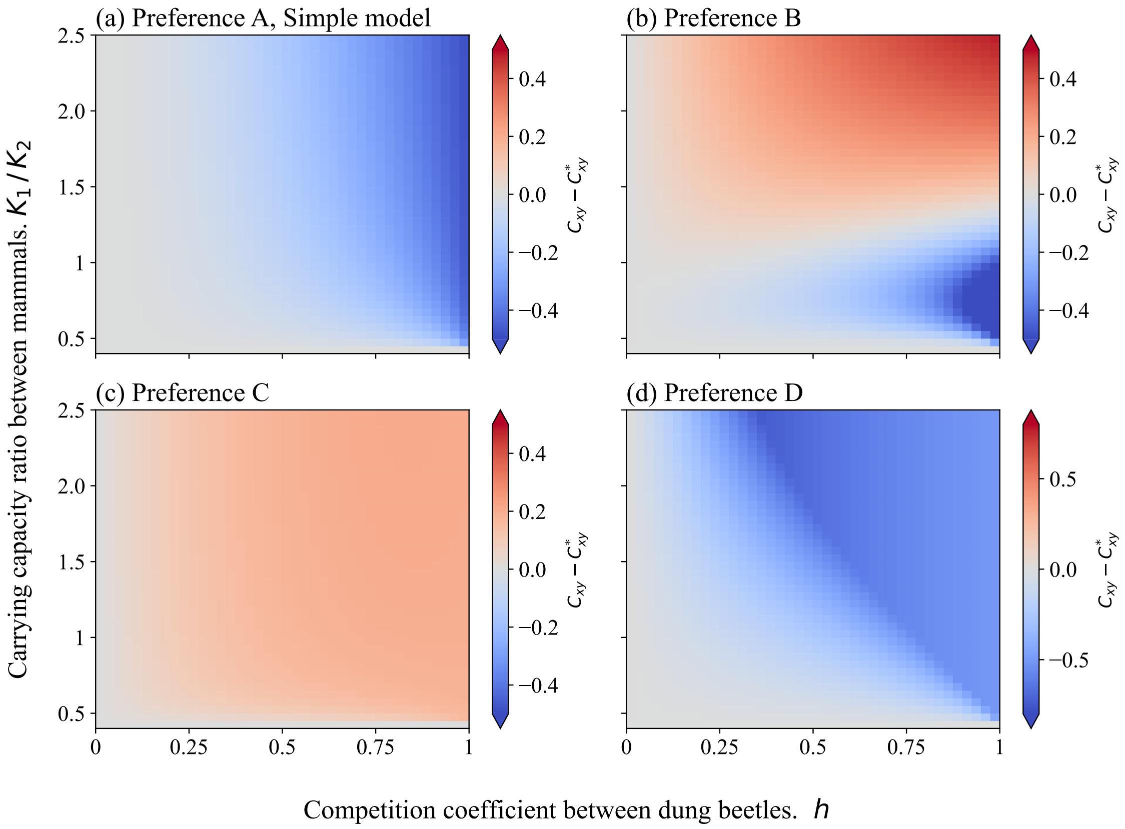


Fig. S11: Interspecific competition strength received by generalsits($C_{xy} = \frac{hG_2y}{kf_2}$) difference from before the invasion. X-axis is the competition coefficient between dung beetles, y-axis When the plot color is red, competition has increased since before the invasion; when the plot color is blue, competition has decreased. Panels (a), (b), (c), and (d) correspond to different fecal preferences of generalist dung beetles: Preference A, Preference B, Preference C, and Preference D, respectively.

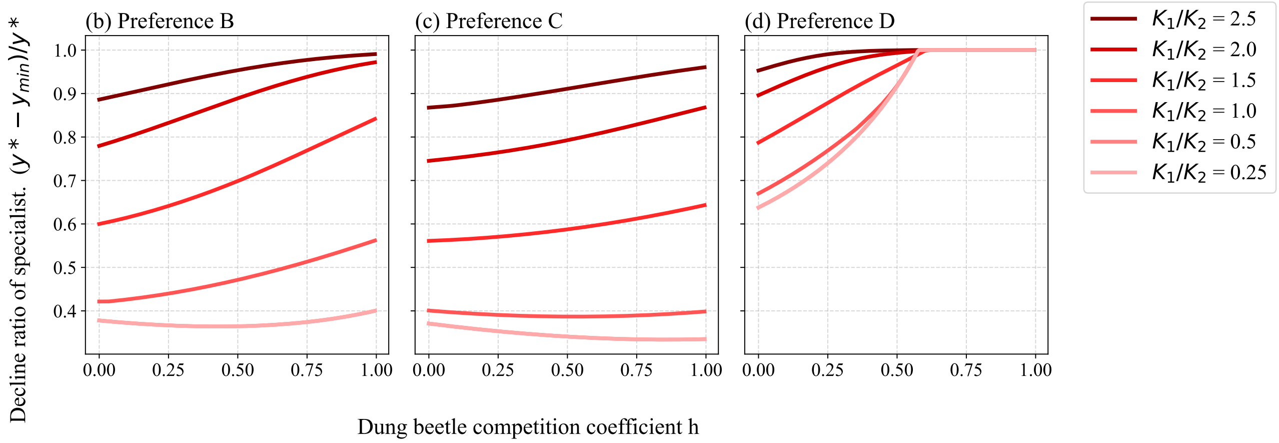


Fig. S12: Decline ratio of specialist dung beetles from before mammal invasion when invasive mammals exhibit irruption dynamics under different fecal preference. The x-axis represents the competition coefficient of dung beetles ($h$), and the y-axis represents the decline ratio of specialists $\left(\frac{y^* - y_{\min}}{y^*}\right)$. Each line represents a different time delay ($\tau$). The fecal preference of generalists is Preference A, and the carrying capacity of invasive mammals is fixed at $K_1 = 1500$Panels (b), (c), and (d) correspond to different fecal preferences of generalist dung beetles: Preference B, Preference C, and Preference D, respectively.
